# Two Comparators May Be All We Need

**DOI:** 10.64898/2026.09.15.751854

**Authors:** Steven M. Muskal

## Abstract

A compound in a cell meets a spectrum of proteins drawn from many families at once, while screening most often interrogates one target at a time. Two questions asked many times in a rank ordering workflow ultimately guide decisions on what gets made and what gets counter-screened, and both are comparative: which of two targets does a compound prefer, and which of two compounds does a target prefer. We built one model for each: the target comparison over a roster of 2,279 human proteins in 34 protein families, the compound comparison over 2,079 in 32. Each model is given two chemical structures and a sequence, or two sequences and a chemical structure, and returns the predicted preference and its prediction strength. No conformational analysis, protein structure, binding site or docked pose is used. The target comparator reached 0.75 accuracy on 8,689 held-out comparisons across families, increasing to 0.94 when the comparisons are filtered by prediction strength. Within one family it reached 0.78 on 32,738 held-out comparisons, increasing to 0.95. The compound comparator reached 0.71 on 65,725 held-out comparisons, increasing to 0.96. Neither compound in any of those comparisons appeared anywhere in training. Accuracy tracks how far apart the two measured potencies actually are. Where the two readings lie within half a log unit of each other the comparators are right 0.57 to 0.64 of the time, and where they differ by more than two log units, 0.90 to 0.92. The compound comparator was also scored one family at a time, where its accuracy ranged from 0.59 to 0.80 across the 31 of its 32 families that had held-out comparisons. Within the chemistry and targets represented in their training data, these models rank compounds and targets. Both models can be explored and downloaded at familyfoundationmodel.com.

## 1. Introduction

A compound in a cell encounters proteins from many families, and its effects reflect its activity across that spectrum[1,2]. Outside proteome-scale chemoproteomic platforms, most screening campaigns ask the question one target at a time, and models have followed. Small molecules are promiscuous by default: chemical similarity alone relates the pharmacology of proteins that share no fold and no family, and off-target activities predicted on that basis are confirmed often enough to reshape a marketed drug’s adverse-effect profile[3,4]. Secondary pharmacology profiling exists for this reason: panels assembled from receptors, channels, transporters and enzymes are run early precisely because activity outside the intended family drives safety-related attrition[5,6]. Within a family the same breadth is measured directly, and kinase inhibitor profiling across the kinome is the standard practice[7,8]. Across families, measurement is uncommon and rarely predicted. Reverse screening, identifying candidate proteins for a given molecule, has gained traction in repurposing[9,10], and large benchmarks establish how well that can be done over enzymes, receptors, channels and proteases together[11,12,13]. Those models score one protein and one compound at a time, so any comparison between two targets is typically recovered afterward by subtracting two separate predictions.

Prioritization in a drug discovery program cycles through the same two comparative operations at every round: which of two targets to counter-screen first, and which of two compounds to make next. Each decision reduces to one of those two. A reverse screen asks which of many potential targets binds a particular compound[10], and a forward screen asks which compound in a batch binds a particular target. Here we compare two targets for a compound or two compounds at a target. In both the answer is a preference, asked over a roster that crosses family boundaries. Comparing two targets against one compound is what medicinal chemistry calls selectivity. A family-level ranking has four immediate uses: (i) prioritizing which family to screen a series against and which counter-screen to run first when no measurement is available; (ii) surfacing an analog series that ranks unexpectedly toward a second family, before an assay is run there; (iii) stating the repurposing question as an ordered target list that can be evaluated over a catalog without choosing a shortlist in advance; and (iv) prioritizing docking by ordering candidate targets from sequence and chemistry before structure-based evaluation. The last use is complementary to pharmacophore retrieval: docking one molecule against 28,579 characterized sites takes 40.5 hours on twenty cores, while retrieval reduces the pool to one that docks in under two minutes[10]. Retrieval selects sites represented in the crystallographic record; the comparator requires no structure.

Three properties favor a direct comparison over predicting an affinity value. First, the same measurements can answer many questions, since every measured compound can be compared with every other one. Second, each comparison uses the same experimental endpoint, so different assay types do not have to be placed on a common scale. This matters because public potency values collected in different formats and laboratories can differ by close to a log unit before modeling begins[14,15]. Finally, the model answers the practical question directly: which compound or target is preferred.

Pairwise formulations are established on the compound axis. PBCNet computes relative binding affinity for two ligands at one receptor from atomic-level structure, and bounded-datapoint classifiers decide which of two molecules is more potent, generally within one target’s data[16,17]. Conditioning that comparison on protein sequence rather than structure is what lets one estimator carry it across a target roster, and sequence-conditioned drug-target models establish that sequence alone supports the underlying task[18,19,20]. The reciprocal question, which of two proteins a compound is more potent at, has been posed within a single family: a kinase comparator introduced both layouts used here[21,22] and a G protein-coupled receptor comparator applies them inside that family[23]. The closest published work on the protein axis trains directly on the difference in activity between two paralogous adenosine receptors and shows that this beats deriving selectivity from two separate affinity models[24]; there the target pair is fixed and the protein is not an input, so the model cannot be asked about a pairing it was not built for. Making the protein an explicit input is what allows one estimator to answer any pairing, and widening the roster past the family boundary is what this paper adds on both axes.

The released target comparator serves 2,279 proteins in 34 families, and the compound comparator serves 2,079 proteins in 32 families, drawn from the liganded human proteome recorded in ChEMBL. ChEMBL 37 defines 40 level-2 protein classes, 36 of which are reachable from a human single protein, and the pool here covers 34 of those 36. The two absent ones are auxiliary channel subunit families whose three human members carry only percentage readings, which have no position on a concentration scale and so cannot enter a comparison at all. At the protein level, 3,213 human single proteins carry a level-2 class, 2,705 of them carry measurements these comparisons can use, and 2,279 are servable by the released target comparator. The question a program usually puts to a single protein is here put to the spectrum a compound actually meets.

This work provides (i) two reciprocal comparators, separately fitted from a public database alone over their respective rosters, with their code and weights released; (ii) their accuracy measured on compound-disjoint holdouts, stratified by family and by the size of the difference being called; and (iii) an assessment of the limited measurement coverage available for cross-family comparisons, determined by the number of compounds measured on both sides of a family boundary.

## 2. Methods

### 2.1 Source data and scope

All data come from ChEMBL 37[25], and no proprietary source enters either model, which is what makes both reproducible and their weights releasable. Records were restricted to human targets of type single protein carrying a level-2 ChEMBL protein class, which supplies the family labels used throughout; those labels are ChEMBL’s own class tree and not a grouping of ours. Accepted endpoints were Ki, IC50, Kd, EC50 and Kb, and rows from PubChem BioAssay were excluded, so no high-throughput screening data contribute. Only exact readings were kept, pActivity was derived from the reported value and unit, not the precomputed pChEMBL column, and repeated readings were reduced to one value per target, compound and endpoint by the median, never by the most potent reading. The pool holds 1,263,626 measurements over 2,705 targets and 794,638 compounds across 34 level-2 families, whose composition is in Supplementary Table S1 and Supplementary Figure S1. Target annotations are UniProt accessions[26]. Units are accepted from a fixed list, M through fM. ChEMBL also carries strings such as 10^2 uM; permissive parsing would have placed three methyltransferases on the wrong scale, so those rows were set aside. The scope accounts for every classified target with no residual, which Supplementary Table S6 traces rule by rule. One pool feeds both comparators, and Table 1 traces it to both holdouts.

**Table 1.** One pool to three holdouts. The same 1,263,626 measurements feed both comparators, and the target comparator forms two kinds of comparison from them: two targets in different families, and two different targets sharing a family. The two arms are fitted together as one model and reported apart throughout, because the held-out set is 79% within-family and a figure pooled over both would follow that mixture rather than either result.

| Step | Target, across families | Target, within one family | Compound comparator |
| --- | --- | --- | --- |
| ChEMBL 37 rows read | 4,636,798 | 4,636,798 | 4,636,798 |
| Measurements in the pool | 1,263,626 | 1,263,626 | 1,263,626 |
| Targets in the pool | 2,705 | 2,705 | 2,705 |
| Compounds in the pool | 794,638 | 794,638 | 794,638 |
| Protein families in the pool | 34 | 34 | 34 |
| Comparisons formed | **89,888** | **333,893** | **2,519,433** |
| Ties dropped | 9,037 | 146,918 | dropped before fitting |
| Fitted comparisons | 81,199 | 301,155 | 1,827,578 |
| Held-out comparisons | **8,689** | **32,738** | **65,725** |
| Discarded as straddling the split | none, one compound per comparison | none, one compound per comparison | 626,130 |
| Compounds fitted, held out, shared | 20,393, 2,195, 0 | 123,177, 13,343, 0 | 474,708, 43,961, 0 |
| Targets appearing in comparisons | 1,867 | 1,987 | 2,079 |
| Family pairings, or families | 292 pairings | 30 families | 32 families |
| First-position win fraction | 0.50 | 0.50 | 0.50 |
| Feature row | 1,998 | 1,998 | 2,556 |

### 2.2 Forming the two comparisons

The target comparison. A comparison exists where one compound carries readings against two targets, under four rules. The endpoint is matched, so a Ki is never ranked against an IC50, and the winner is the target carrying the higher pActivity on that endpoint; the comparison therefore predicts the direction of the difference between the two readings and not its magnitude, so a selectivity window is not an output, only its sign. The two targets are different proteins, which matters because 45 human proteins in ChEMBL carry two level-2 labels, most often reader and writer, and a single protein filed twice would otherwise present itself as a comparison. Each comparison is then filed by whether its two targets share a family label, giving two arms that are fitted as one model and reported separately: across families, where the two labels are disjoint, and within one family, where two different proteins share one. Ties were dropped, since two equal readings carry no answer. And which target occupies the first position was randomized per comparison, so no model profits from guessing a slot. Comparisons were capped at 50 per compound and 20,000 per family pairing and endpoint, each cap applied within an arm rather than across both, giving 89,888 comparisons across families over 1,867 targets and 292 family pairings, and 333,893 within a family over 1,987 targets in 30 families.

The compound comparison. The reciprocal comparison exchanges the roles, and it is a separately fitted model with its own split and its own holdout. It exists wherever two compounds were measured at one target on the same endpoint. The winner is the compound carrying the higher pActivity, ties are dropped before fitting, and the first position is randomized, so the label balance is exactly 0.50.

Both compounds in a comparison are measured at the same protein, so any single comparison stays within one protein family. The model itself is not limited to one family, since it can be asked about proteins from all 34. The same model handles a receptor, a channel or an acetyltransferase without any change. The two comparisons reach that breadth differently: a target comparison sets two proteins against each other, either across families or within one, while a compound comparison asks about one protein at a time and can ask about any of them. This is the layout introduced for kinases[21] on a roster no longer confined to one family.

Preference is exact here because every comparison is endpoint matched and every reading is converted to pActivity, the negative logarithm of a concentration. Higher is therefore preferred uniformly across all five endpoints, with no direction to flip between an inhibition endpoint and an agonist one: both readings come from the same target and the same endpoint, and the model predicts which of the two compounds is active at the lower concentration there.

The same measurements yield 2,519,433 compound comparisons and 423,781 target comparisons, and the target comparisons are themselves unevenly split, 89,888 across families against 333,893 within one. A target comparison across families needs one compound measured at two targets in different families, and only 4.6% of compounds are; a compound comparison needs two compounds at the same target, where each further compound tested can be compared with every one already measured. The difference is in the record, not in the method.

### 2.3 Features, model and augmentation

Two feature blocks are defined once and serve both comparators. A sequence becomes the 480-dimensional residue-wise mean of ESM2-t12-35M over the full canonical sequence, with terminal tokens excluded[27,28]. A compound becomes a 1,024-value radius-2 Morgan count fingerprint, which stores substructure counts, not presence, plus 14 whole-molecule descriptors: molecular weight, heavy-atom count, bond count, rotatable-bond count, ring count, and counts of C, N, O, S, F, Cl, Br, I and P[29]. Both blocks are byte-identical between the two comparators, verified directly.

Each row places the two things being compared on the outside and what they share in the middle, so the order is the question (Figure 1). The target row is sequence A, compound, sequence B, for 1,998 values; the compound row is compound A, sequence, compound B, for 2,556. No structure, pocket, pose or contact map is used. Both estimators are conventional random forests[30,31], and both use 300 trees, square-root feature sampling, bootstrap resampling, the Gini criterion, no depth limit and seed 0. They differ only in minimum leaf size, 8 for the target comparator and 20 for the compound comparator. Both return a probability and never an affinity.

**Figure 1.**
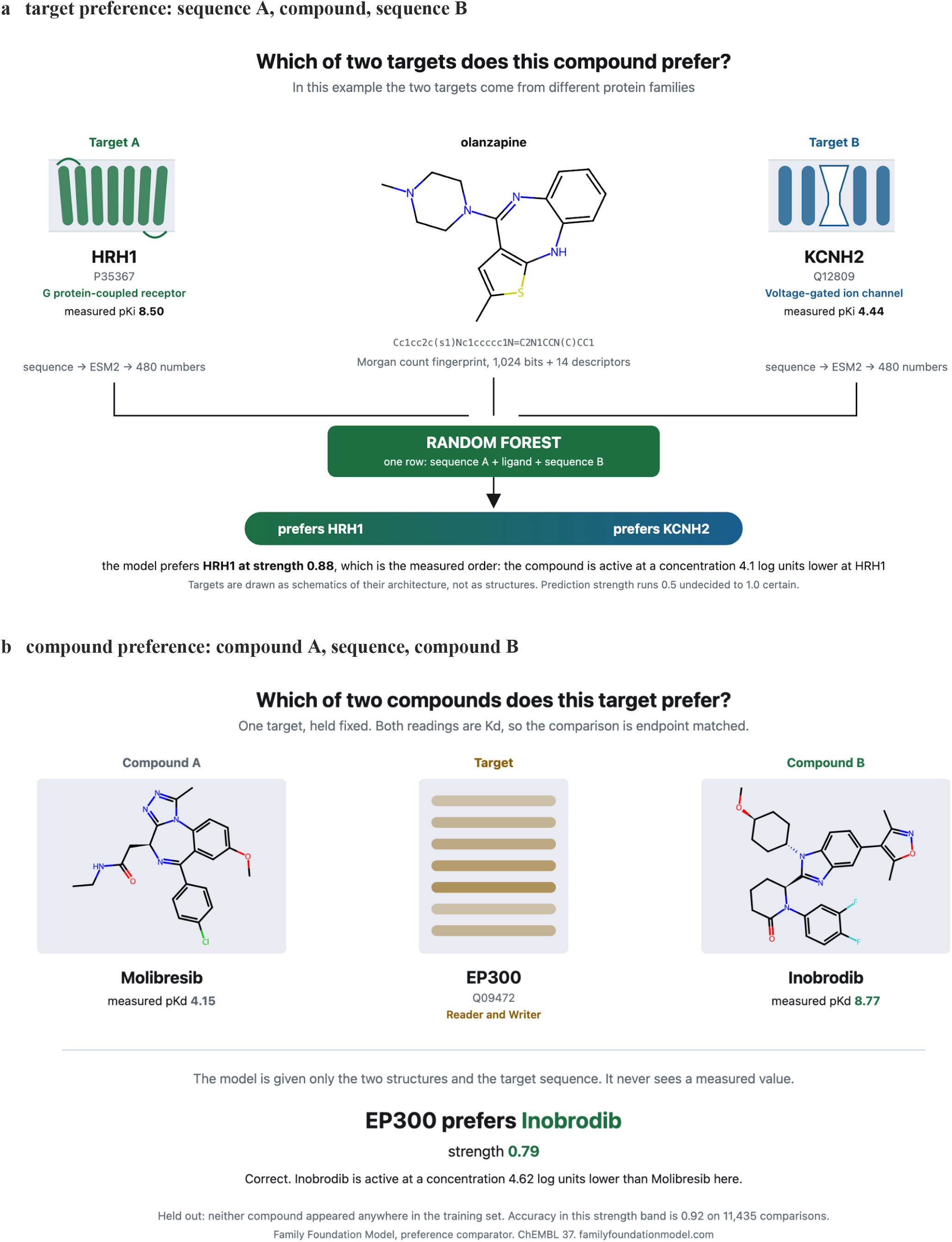
The two comparisons, each with a real case. Each comparison is one feature row, and the order is the question: the two things compared take the outer positions and what they are compared against sits between them. Every sequence enters as 480 ESM2-t12-35M numbers, the mean over residues of the full canonical sequence, and every compound as a 1,024-value radius-2 Morgan count fingerprint plus 14 whole-molecule descriptors, and the two blocks are byte-identical between the panels. Each forest returns one probability, drawn as the bar, and prediction strength runs from 0.50 undecided to 1.00 certain. Panel a, the target comparison, is sequence A, compound, sequence B, for 1,998 values; the two targets in this example carry disjoint level-2 ChEMBL family labels, so this comparison reaches across a family boundary. The case is olanzapine put to the histamine H1 receptor (HRH1) and to hERG (KCNH2), measured at pKi 8.50 and 4.44, where the model answers 0.88 for H1, the measured order; both readings are in the training data, so this case shows what the model returns, not a test of it. Panel b, the compound comparison, is compound A, sequence, compound B, for 2,556 values, anchored at one target, so the family span is carried by the roster the model serves. The case is held out, and neither compound appeared anywhere in fitting: the acetyltransferase p300 (EP300) against molibresib and inobrodib, both clinical-stage bromodomain inhibitors, measured at pKd 4.15 and 8.77, where the model answers 0.79 for inobrodib, the measured order, in a strength band whose accuracy is 0.92 on 11,435 comparisons. Targets are drawn as schematics of their architecture, not as structures, and no structure is a model input.

Random forests kept fitting inexpensive while allowing the released models to be extended and checked exactly. The target comparator fitted 764,708 rows by 1,998 columns in 239 s, and the compound comparator fitted 3,655,156 rows by 2,556 columns in 1,257 s; each required one CPU pass, no GPU and no warm start. New trees can be fitted only on a contributor’s measurements and concatenated with the released 300 trees, without changing those trees or requiring the original training corpus; the new trees receive the bounded vote share k/(300+k), which the utilities measure on a reference set spanning the target roster. A worked example shows what adding data buys. Eighty trees were fitted on 1,858 comparisons a user held and the released model did not, across five targets, and tested against 797 further comparisons from those same targets that were held back. Accuracy on those five targets rose from 0.69 to 0.73, with no measured loss across 1,500 reference comparisons spanning 967 targets. Fitting the same 80 trees on measurements the released model already held raised accuracy only to 0.70, so most of the gain comes from measurements the model has not seen. Each bundle uses seed 0, and installation succeeds only when 30 reference predictions reproduce to within 1e-6.

Every comparison is entered twice, with the outer blocks exchanged and the label inverted. The reversed row adds no experimental information, but the row is positional, so the two orientations are distinct points in the input space and the expected symmetry enters as supervised information. Predictions average both orders. Three exact consequences were verified on the target comparator: the two orders sum to one, a target against itself returns 0.50, and the fitted label balance is 0.50. On the compound comparator the label balance is likewise exactly 0.50, and a compound against itself returns exactly 0.50.

The two target arms are one model: 382,354 fitted comparisons over 135,852 compounds, 41,427 held out over 14,719, and a roster of 2,279 servable accessions.

### 2.4 Split and evaluation

Neither model was tested on a compound it had been fitted on. The two splits get there differently. Compounds were assigned by a BLAKE2b digest of their InChIKey. The target comparison carries one compound, so withholding a compound withholds every comparison it appears in. The compound comparison carries two, so compound-disjoint has to mean both: a comparison is fitted only if both compounds are fit-side and held out only if both are test-side, and the 626,130 comparisons of 2,519,433 that straddle the split are discarded. That cost is what makes the holdout strict. Neither split shares a compound between its sides, verified on every run, and Table 1 gives both in full.

A comparison is held out only when both of its compounds are, so withholding 20% of the compounds sets aside only about 4% of the comparisons. An unstratified split holds out 0.04 of comparisons in every family, but family sizes differ 600-fold and the smallest contributes fewer than 200 held-out comparisons. The released model therefore sets the fraction per family, solving for the square root of a target count over the family’s size and clipping to a floor and a ceiling, so each family is aimed at a comparable absolute count, not the same share. That is what makes the per-family figures in 3.4 comparable and allows a table by family.

Accuracy is the share of held-out comparisons whose winner a model names correctly, and it is the only performance measure reported. Prediction strength is the larger of the two output probabilities and runs from 0.50 to 1.00. Since every comparison is entered in both orders, the two possible answers are exactly balanced in fitting and in evaluation, so neither figure can be inflated by an uneven split.

## 3. Results

### 3.1 Ranking two targets

The target comparator picks the target with the higher measured activity 0.75 of the time where the two targets come from different families, over 8,689 comparisons drawn from 2,195 compounds the model never saw in fitting, spanning 180 of the 287 family pairings on its fitted side. Asked about two different targets of one family, it is correct 0.78 of the time over 32,738 comparisons from 13,343 unseen compounds, across 30 families.

Within-family accuracy runs from 0.69 to 0.94 across the 17 families holding 200 or more held-out comparisons (Supplementary Table S8). Kinase is the hardest of them at 0.73 on 4,979 comparisons over 360 proteins, despite holding the most data of any family, and Family A G protein-coupled receptor reads 0.76 on 5,412. For a reference point, the G protein-coupled receptor comparator answers the same question inside that single family at 0.80 on its own unseen compounds[23].

### 3.2 Ranking two compounds

The compound comparator picks the compound a target prefers 0.71 of the time, over 65,725 comparisons in which neither compound appeared in fitting. A further 626,130 comparisons had one compound on each side of the split and were discarded rather than assigned to either, which is what makes the test a strict one.

### 3.3 Prediction strength

Prediction strength is the larger of the two returned probabilities, and in both models accuracy rises with it while coverage falls (Figure 2, Table 2). At strength 0.80 the target comparator answers 2,980 of its 8,689 across-family comparisons at 0.94 and 12,529 of its 32,738 within-family comparisons at 0.95, and the compound comparator 3,995 of its 65,725 at 0.96. The relationship holds band by band and not only cumulatively, from 0.55 among the target comparator’s weakest across-family calls to 0.97 among its strongest, and from 0.52 to 0.98 within a family, so the returned probability carries information beyond the winner it names. Strength, rather than the headline, is what a user acts on.

**Table 2.** The comparisons side by side, each measured on its own held-out set of 8,689, 32,738 and 65,725 comparisons. An accuracy at a given prediction strength is measured only on the comparisons that reach it, and the share answered gives how many those are. The first two columns come from one model asked two kinds of question. The last two rows give accuracy where the two measurements being compared are closest together and furthest apart.

|  | Target, across families | Target, within one family | Compound preference, two compounds |
| --- | --- | --- | --- |
| **all held-out comparisons** | **0.75** | **0.78** | **0.71** |
| prediction strength 0.60, share answered | 0.82, 0.71 | 0.85, 0.76 | 0.84, 0.43 |
| prediction strength 0.70, share answered | 0.89, 0.49 | 0.91, 0.56 | 0.92, 0.17 |
| prediction strength 0.80, share answered | 0.94, 0.34 | 0.95, 0.38 | 0.96, 0.06 |
| prediction strength 0.90, share answered | 0.97, 0.21 | 0.98, 0.21 | 0.99, 0.01 |
| within half a log of true difference | 0.59 on 2,264 | 0.64 on 11,228 | 0.57 on 20,200 |
| beyond two logs of true difference | 0.91 on 2,288 | 0.92 on 5,254 | 0.90 on 11,673 |

**Figure 2.**
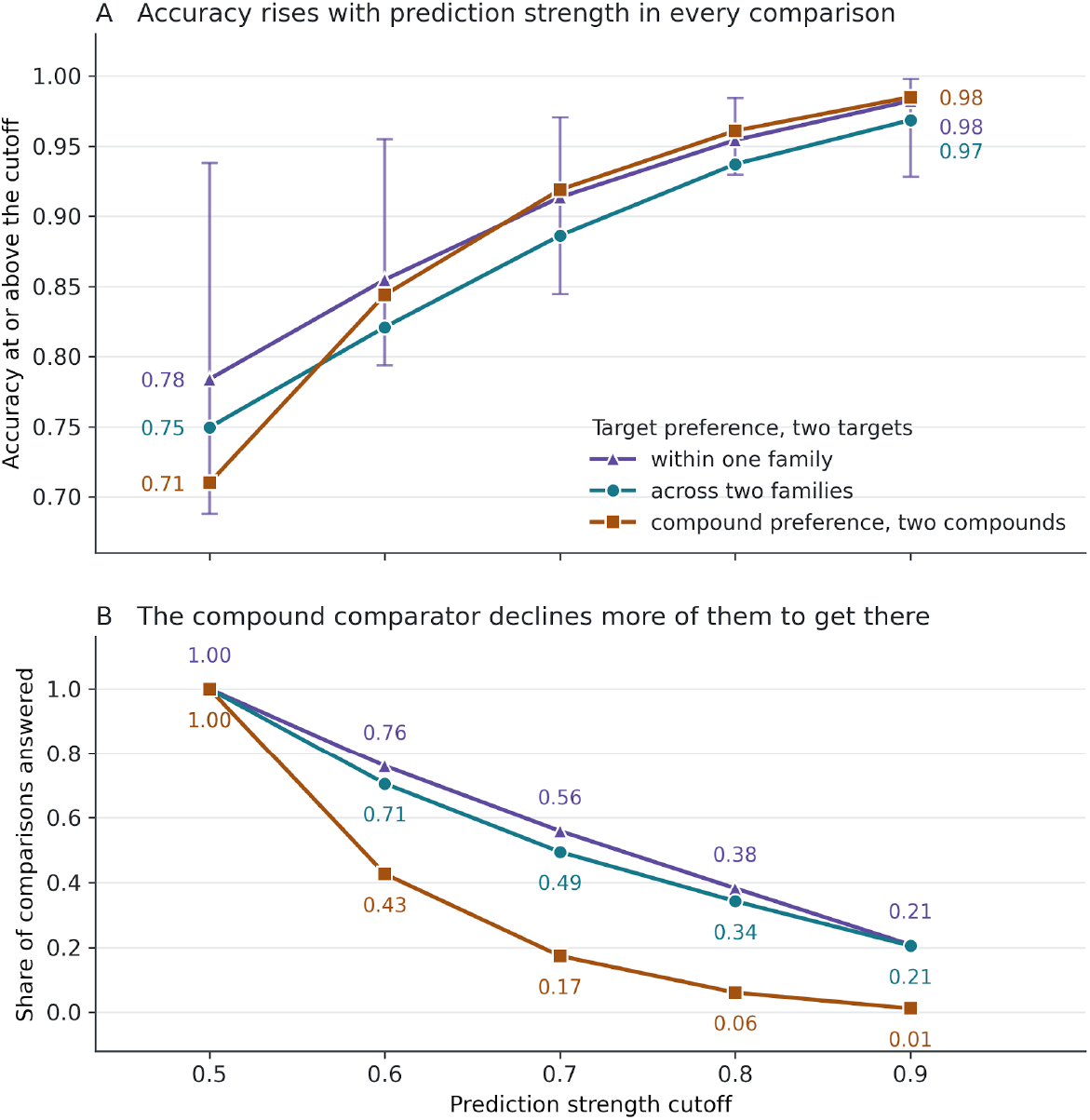
Accuracy against prediction strength, all three comparisons on one axis. Prediction strength is the larger of the two returned probabilities and runs from 0.50 to 1.00. Panel A gives accuracy over the held-out comparisons at or above each cutoff and panel B the share of each holdout surviving it, over 8,689 target comparisons across two families, 32,738 target comparisons within one family, and 65,725 compound comparisons. Vertical bars on the within-family series give the range across the 17 families carrying 200 or more held-out within-family comparisons at that cutoff, from 0.69 to 0.94 ungated and from 0.93 to 1.00 at strength 0.90, so the curve is a position within a spread rather than a single family’s result. The within-family series sits above the across-family series at every cutoff while answering a similar share, and the compound comparator reaches the highest accuracy of the three above 0.60 by declining far more comparisons to get there, keeping 0.06 at strength 0.80 against 0.34 and 0.38 for the two target series.

### 3.4 What governs a call

The size of the real difference governs both comparators the same way. Where the two readings sit within half a log accuracy is 0.59 on 2,264 across-family target comparisons, 0.64 on 11,228 within-family ones and 0.57 on 20,200 compound comparisons, and beyond two logs it is 0.91,0.92 and 0.90 respectively (Table 2). While a near-tie is where a comparator is least useful, it is also where the two measurements are least separable to begin with, since inter-assay variability is largest there.

For the target comparison across families the second factor is which two families are being asked about. Among pairings carrying 100 or more held-out comparisons, kinase against voltage-gated ion channel reads 0.94 on 301 and cytochrome P450 against kinase 0.89 on 561, while electrochemical transporter against Family A G protein-coupled receptor reads 0.68 on 1,016. That last pairing is the aminergic case, where monoamine transporters and aminergic receptors share their ligands, and it is among the pairings the comparator separates less well (Supplementary Table S2).

For the compound comparison the second factor is the roster itself. Accuracy runs from 0.59 to 0.80 across the 31 families carrying held-out comparisons, 27 of which hold out 500 or more, and the kinase family reads 0.69 on 9,585, below the pooled figure rather than above it (Supplementary Figure S2, Supplementary Table S3). A single pooled accuracy would conceal that spread, which is why the holdout fraction is set per family rather than globally.

The endpoint mix separates the two comparators. The compound comparison holds 9,375 EC50 and 538 Kb comparisons, reading 0.71 and 0.73, so functional and agonist readings are within its reach; the target comparison holds 945 EC50 comparisons across families, of which 114 fall in the holdout, and no Kb at all across families, since it needs the same endpoint measured at two targets in different families and agonist data are rarely collected that way (Supplementary Table S4). The within-family target holdout also carries 2,815 EC50 and 73 Kb comparisons, at 0.81 and 0.78, respectively.

### 3.5 Cross-family coverage sets the across-family ceiling

A target comparison across two families requires one compound measured on both sides of a family boundary, and almost no compound is. Inside a family the record is far deeper: the same pool yields 1,643,762 comparisons between two targets of one family against 518,194 across a boundary, and 69% of that within-family volume is kinase against kinase. Of 785,745 compounds carrying a classified human medicinal-chemistry measurement, 749,812 are measured in a single family; 33,158 are measured in two, 2,115 in three, and 660 in four or more (Figure 3). Only 4.6% cross a boundary at all, and the median compound that does crosses exactly one. Measurement that deepens one family cannot form a comparison that needs two, so the ceiling here is set by the published record rather than by the model.

**Figure 3.**
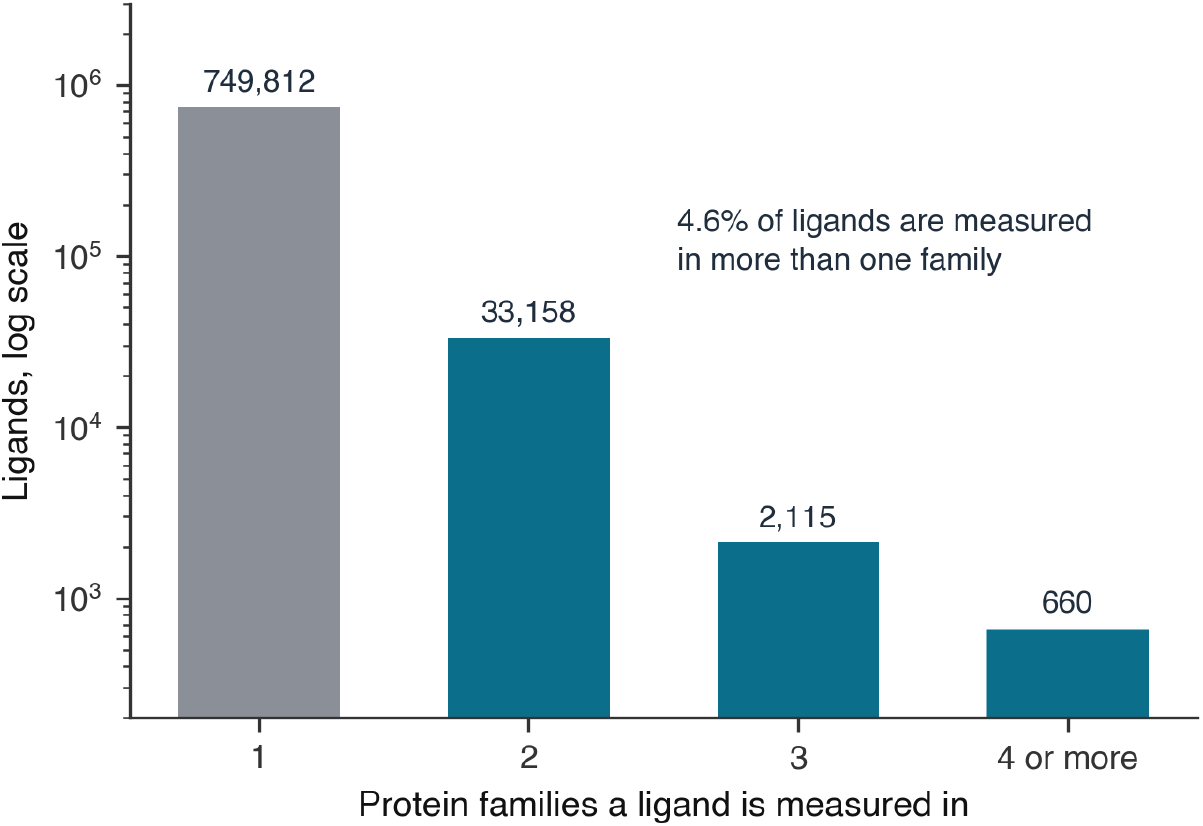
Why the target-comparison ceiling is where it is. The number of compounds measured in one, two, three, or four or more level-2 protein families, over the 785,745 carrying a classified human medicinal-chemistry measurement, on a logarithmic count axis. A target comparison requires a compound measured on both sides of a family boundary, and 749,812 of these are measured inside a single family. Neither the within-family target comparison nor the compound comparison is bounded this way, since both draw on compounds measured inside a single family, which is why the same pool yields 333,893 and 2,519,433 of them against 89,888. The counts describe the available record and not how promiscuous compounds are, since a compound measured in one family may never have been tested in another.

## 4. Discussion

### 4.1 What the model supports

We trained two reciprocal models: one compares targets for a compound, the other compares compounds at a target. The target comparator serves 2,279 proteins in 34 families and the compound comparator serves 2,079 proteins in 32 families; both predict relative preference. They run from sequence and two-dimensional chemistry alone, with no structure, pocket or pose; one comparison is a forest evaluation, so a panel or list can be ordered outright. PharmCast takes the same structure-free stance on a different descriptor, predicting a three-dimensional pharmacophore fingerprint directly from two-dimensional structure without conformer generation; that fingerprint also feeds the reverse screen[10,32].

Either comparator can be extended on a user’s own machine, as 2.3 describes, and an extended model carries no performance figure from this paper until it is measured on the extender’s own held-out comparisons.

The kinase and G protein-coupled receptor foundation models are drill-downs in this sense: the Family Foundation Model establishes breadth across the liganded proteome, and the family models show the depth a focused corpus can add. Within-family accuracy is where a focused corpus has most to add, and it is lowest in the family that already holds the most data: kinase reads 0.73 against 0.85 for transferase and 0.94 for the Toll-like and Il-1 receptors, because kinases resemble one another and volume does not answer that. Together they are worked examples of the same breadth-to-depth choice for a user with proprietary measurements. The kinase model is that case in full: it is fitted on a curated corpus outside ChEMBL and checked against ChEMBL, which is the arrangement a contributor with their own measurements is in.

The split is what the compound-comparison figure rests on: both compounds unseen, and every straddling comparison discarded rather than assigned. Panel b of Figure 1 is one such case: p300 against molibresib and inobrodib, two clinical-stage bromodomain inhibitors, so the discrimination is between drugs of one mechanistic class, in a family neither sibling model covers.

Panel a is the target comparison in one picture. Olanzapine is put to the histamine H1 receptor and to hERG, proteins that share no fold, no family and no assay convention, which is exactly the comparison a program asks when an aminergic drug has a channel liability. Both readings there are in the training data, so the panel illustrates the output a user acts on, a named winner with a strength attached, on a pairing no within-family model can be asked about at all.

Given a feature row built around one shared element, exactly two pairwise questions are available: hold the target and pair the compounds, or hold the compound and pair the targets. The pair is the complete set rather than a selection from a larger one, which is why two comparators may be all the ranking a prioritization cascade requires. These rankings help users prioritize the next experimental step alongside measurement; both models return relative preference without estimating absolute potency.

Users can select a prediction-strength threshold to trade coverage for higher observed accuracy, as shown in Figure 2. Cost and coverage here are operating properties of the released comparators. Crossing a family boundary costs three points, 0.75 against 0.78, measured on one model asked both questions. For reference, the in-family G protein-coupled receptor target comparator reads 0.80 on its own compound-disjoint holdout[23], against 0.76 for this model inside that same family; the two sibling within-family compound comparators bracket the compound-comparison result on splits that are not comparable to it (Supplementary Table S5).

### 4.2 What limits it

For the target comparison across families, accuracy tracks the family pairing more than the chemistry. Within a family, target preference varies with compound chemistry, so unusual compounds are difficult cases. For the compound comparison the reverse holds, and a target whose preferences depart from the usual ordering of the two compounds is the hard case.

Near-ties are difficult for both models: accuracy runs 0.57 to 0.64 where the two measurements are within half a log unit, and 20,200 of the compound comparator’s 65,725 held-out comparisons fall inside that band. Raising the prediction-strength threshold reduces coverage, and decisions that depend on distinguishing near-ties require measurement.

The aminergic pairing is a well-populated but difficult case, at 0.68 on 1,016 comparisons, and one where a cross-family ranking would be most valuable, since monoamine transporters and aminergic receptors are where repurposing and off-target liability meet.

Cross-family comparisons are limited by the available measurement coverage: only 4.6% of classified compounds are measured at targets in different families (Figure 3). Within a family, measurements are more numerous, while similarity between proteins makes their preferences harder to distinguish.

Every figure here was measured on chemistry and targets from the distribution the models were fitted on. Withholding compounds is stricter than withholding individual comparisons and weaker than withholding whole targets, and how strictly a split is drawn is itself a major determinant of reported performance[33,34,35]. Results therefore apply to represented families and targets; a target outside the served roster is refused, never guessed at.

Both models return a probability, so there is no predicted affinity here to report as one. Nothing measured in this work connects cross-family breadth to toxicity. Breadth recorded in a public database partly records how many assay panels a compound was put through, and testing that link would require a real safety outcome set.

### 4.3 Using it to prioritize

The two comparators can be used in sequence to help users prioritize different tasks: the compound comparator ranks candidates to make or buy, and the target comparator ranks targets for counter-screening. For a compound of interest, users can compare its intended target with each candidate target and prioritize candidates with higher predicted preference. At a prediction-strength threshold of 0.80, target-comparison accuracy is 0.94 across families with coverage of 0.34, and 0.95 within one family with coverage of 0.38. A prediction below the selected prediction-strength threshold leaves the comparison unanswered and gives no basis for inferring similar activity. At the same threshold, compound-comparison accuracy is 0.96 with coverage of 0.06, allowing users to prioritize the retained comparisons while leaving the remaining comparisons unanswered. Targets are named by UniProt accession throughout, because gene symbols are many-to-many against accessions and the failure is silent.

## 5. Conclusion

Two reciprocal comparators answer the two questions a program asks, the target comparator over 2,279 proteins in 34 families and the compound comparator over 2,079 in 32: given a compound and two proteins, which protein is preferred, and given a target and two compounds, which compound is preferred. The target comparator reaches 0.75 on 8,689 compound-disjoint comparisons between two families and 0.78 on 32,738 between two proteins of one family, and the compound comparator 0.71 on 65,725 in which neither compound was fitted. Accuracy rises with the strength of the answer and the measured difference, and the compound result spans 0.59 to 0.80 across 31 families. Within represented chemistry and targets, these released models rank compounds and targets; they do not estimate potency.

The across-family target comparison is limited by the published record. Only 4.6% of classified compounds are measured in more than one protein family, and those few usually reach only two, so more measurements inside the family a compound was designed against cannot raise this ceiling. What raises it is compounds measured against targets in other families, with the results published. No single organization can cover that space, but a released model of this kind improves for everyone as the record fills, across the many protein families that disease requires us to consider.

## 6. Data, Code, and Model Availability

Both models are trained on ChEMBL 37 and on nothing else, so no commercial corpus restricts their release. The code is Apache 2.0 and the weights are CC BY-SA 4.0. The models are downloadable at familyfoundationmodel.com[36], which also enables interactive experimentation and carries the target roster, methods and measured limitations, and the tools to leverage them are downloadable at github.com/smuskal/ffm[37]. Installation is git clone followed by install.sh, which pins the environment, fetches the bundle and admits the installation only when 30 reference predictions reproduce within 1e-6.

User measurements enter the same local pairwise workflow, so confidential data remain on the user’s machine.

Both run locally on CPU and send nothing elsewhere. The compound comparator is fitted on 4.8 times as many rows and carries 46,926,130 tree nodes against 19,713,090. The target comparator is 0.42 GB on disk, about 3.4 GB resident, loads in about 5 s and serves 2,279 targets; an 8 GB laptop runs it comfortably. The compound comparator is 1.18 GB on disk, about 5.4 GB resident, reaches 7.8 GB while unpickling, loads in about 11 s and serves 2,079 targets; it needs more than 8 GB of memory. One comparison takes 0.11 s and a batch of 1,000 takes 0.07 s, so loading dominates cost. The pinned environment is Python 3.10.15, scikit-learn 1.7.2, numpy 2.2.6, RDKit 2025.09.5, torch 2.9.1 and transformers 4.57.3; forests require the released scikit-learn version to load.

The sibling models are also available: the kinase comparators at kinasefoundationmodel.com[22] and the G protein-coupled receptor comparator at gpcrfoundationmodel.com [23].

## 7. Author Contributions

S.M.M. conceived the comparison tasks, assembled the source data, built and evaluated the models, and wrote the manuscript.

## 9. Acknowledgments

This work uses ESM2, RDKit, scikit-learn, ChEMBL and UniProt.

## 10. Competing Interests

S.M.M. is founder and CEO of Eidogen-Sertanty, Inc., which develops the models described here.

Large language model agents were used for code generation, experiment execution and internal audit under human direction. No language model is an author, and every scientific claim here is the author’s own.

