## Supplementary Material: Tables S1 to S9 and Figures S1 and S2 for "Two Comparators May Be All We Need"

Supplementary Table S1. The training pool by protein family. Measurements are the ChEMBL 37 pool of 1,263,626 readings, counted per level-2 family. A target carrying two family labels is counted in both, so the measurement column sums to 1,272,042, not the pool total. Targets are the roster the released target comparator can score, taken from the model release. Thirty-two roster entries carry two labels, most often reader with writer. A pair of targets sharing either label is a within-family comparison rather than an across-family one, which is what keeps a single protein filed twice from presenting itself as a pair of families. The distribution is what sets which pairings carry enough comparisons to measure.

| Family | Measurements | Targets |
|---|---:|---:|
| Kinase | 325,634 | 422 |
| Family A G protein-coupled receptor | 281,585 | 205 |
| Protease | 119,156 | 191 |
| Oxidoreductase | 59,784 | 191 |
| Transferase | 57,786 | 268 |
| Nuclear receptor | 55,539 | 39 |
| Hydrolase | 42,595 | 265 |
| Voltage-gated ion channel | 40,423 | 78 |
| Eraser | 38,872 | 37 |
| Lyase | 37,059 | 30 |
| Cytochrome P450 | 28,509 | 33 |
| Electrochemical transporter | 27,684 | 89 |
| Reader | 26,206 | 81 |
| Phosphodiesterase | 22,685 | 26 |
| Ligand-gated ion channel | 19,438 | 50 |
| Writer | 13,697 | 37 |
| Phosphatase | 13,644 | 81 |
| Toll-like and Il-1 receptors | 13,501 | 7 |
| Family C G protein-coupled receptor | 9,000 | 10 |
| Other ion channel | 8,603 | 19 |
| Family B G protein-coupled receptor | 7,473 | 15 |
| Primary active transporter | 6,207 | 23 |
| Isomerase | 5,736 | 31 |
| Ligase | 5,127 | 30 |
| Aminoacyltransferase | 2,174 | 6 |
| Fatty acid binding protein family | 1,381 | 6 |
| Frizzled family G protein-coupled receptor | 750 | 4 |
| Calcium channel auxiliary subunit alpha2delta family | 653 | 2 |
| Taste family G protein-coupled receptor | 554 | 25 |

| Family | Measurements | Targets |
|---|---|---|
| Transmembrane 1-electron transfer carriers | 319 | 5 |
| Group translocator | 261 | 2 |
| Slow voltage-gated potassium channel accessory protein family | 5 | 1 |
| Calcium-activated potassium channel auxiliary subunit slowpoke-beta family | 1 | 1 |
| Sodium channel auxiliary subunit beta family | 1 | 1 |

Supplementary Table S2. Target-comparison accuracy by family pairing. Every pairing carrying 30 or more held-out across-family target comparisons, measured on the 8,689-comparison holdout. A pairing's accuracy reflects which targets and which chemistry happen to have been measured on both sides of that boundary, so it is not a statement about how separable two families are in general. Pairings below 30 are omitted as too thin to read.

| Family pairing | n | Accuracy |
|---|---|---|
| Eraser, Phosphodiesterase | 41 | 0.95 |
| Kinase, Voltage-gated ion channel | 301 | 0.94 |
| Phosphodiesterase, Voltage-gated ion channel | 31 | 0.94 |
| Cytochrome P450, Transferase | 78 | 0.91 |
| Protease, Voltage-gated ion channel | 72 | 0.90 |
| Cytochrome P450, Kinase | 561 | 0.89 |
| Electrochemical transporter, Voltage-gated ion channel | 87 | 0.89 |
| Family A G protein-coupled receptor, Primary active transporter | 50 | 0.88 |
| Eraser, Voltage-gated ion channel | 33 | 0.88 |
| Cytochrome P450, Nuclear receptor | 56 | 0.88 |
| Kinase, Primary active transporter | 31 | 0.87 |
| Kinase, Reader | 64 | 0.86 |
| Cytochrome P450, Protease | 99 | 0.85 |
| Family A G protein-coupled receptor, Voltage-gated ion channel | 371 | 0.84 |
| Nuclear receptor, Voltage-gated ion channel | 34 | 0.82 |
| Cytochrome P450, Electrochemical transporter | 102 | 0.82 |
| Cytochrome P450, Primary active transporter | 61 | 0.82 |
| Family A G protein-coupled receptor, Writer | 33 | 0.82 |
| Hydrolase, Protease | 114 | 0.81 |
| Electrochemical transporter, Kinase | 36 | 0.81 |
| Cytochrome P450, Oxidoreductase | 191 | 0.80 |
| Oxidoreductase, Voltage-gated ion channel | 54 | 0.80 |
| Kinase, Protease | 38 | 0.79 |
| Family A G protein-coupled receptor, Oxidoreductase | 64 | 0.78 |

| Family pairing | n | Accuracy |
|---|---:|---:|
| Eraser, Oxidoreductase | 154 | 0.78 |
| Cytochrome P450, Family A G protein-coupled receptor | 364 | 0.76 |
| Hydrolase, Lyase | 38 | 0.76 |
| Eraser, Transferase | 84 | 0.75 |
| Kinase, Transferase | 1,544 | 0.74 |
| Oxidoreductase, Protease | 31 | 0.74 |
| Family A G protein-coupled receptor, Nuclear receptor | 49 | 0.73 |
| Cytochrome P450, Eraser | 64 | 0.73 |
| Cytochrome P450, Voltage-gated ion channel | 256 | 0.73 |
| Kinase, Oxidoreductase | 140 | 0.73 |
| Hydrolase, Oxidoreductase | 83 | 0.72 |
| Group translocator, Transferase | 30 | 0.70 |
| Lyase, Protease | 30 | 0.70 |
| Eraser, Kinase | 238 | 0.69 |
| Cytochrome P450, Hydrolase | 45 | 0.69 |
| Electrochemical transporter, Family A G protein-coupled receptor | 1,016 | 0.68 |
| Family A G protein-coupled receptor, Kinase | 131 | 0.64 |
| Electrochemical transporter, Primary active transporter | 119 | 0.64 |
| Family A G protein-coupled receptor, Ligand-gated ion channel | 118 | 0.60 |
| Electrochemical transporter, Protease | 59 | 0.59 |
| Hydrolase, Kinase | 171 | 0.58 |
| Family A G protein-coupled receptor, Hydrolase | 53 | 0.57 |
| Eraser, Reader | 59 | 0.53 |
| Kinase, Lyase | 32 | 0.50 |
| Family A G protein-coupled receptor, Family B G protein-coupled receptor | 54 | 0.50 |
| Isomerase, Kinase | 56 | 0.21 |

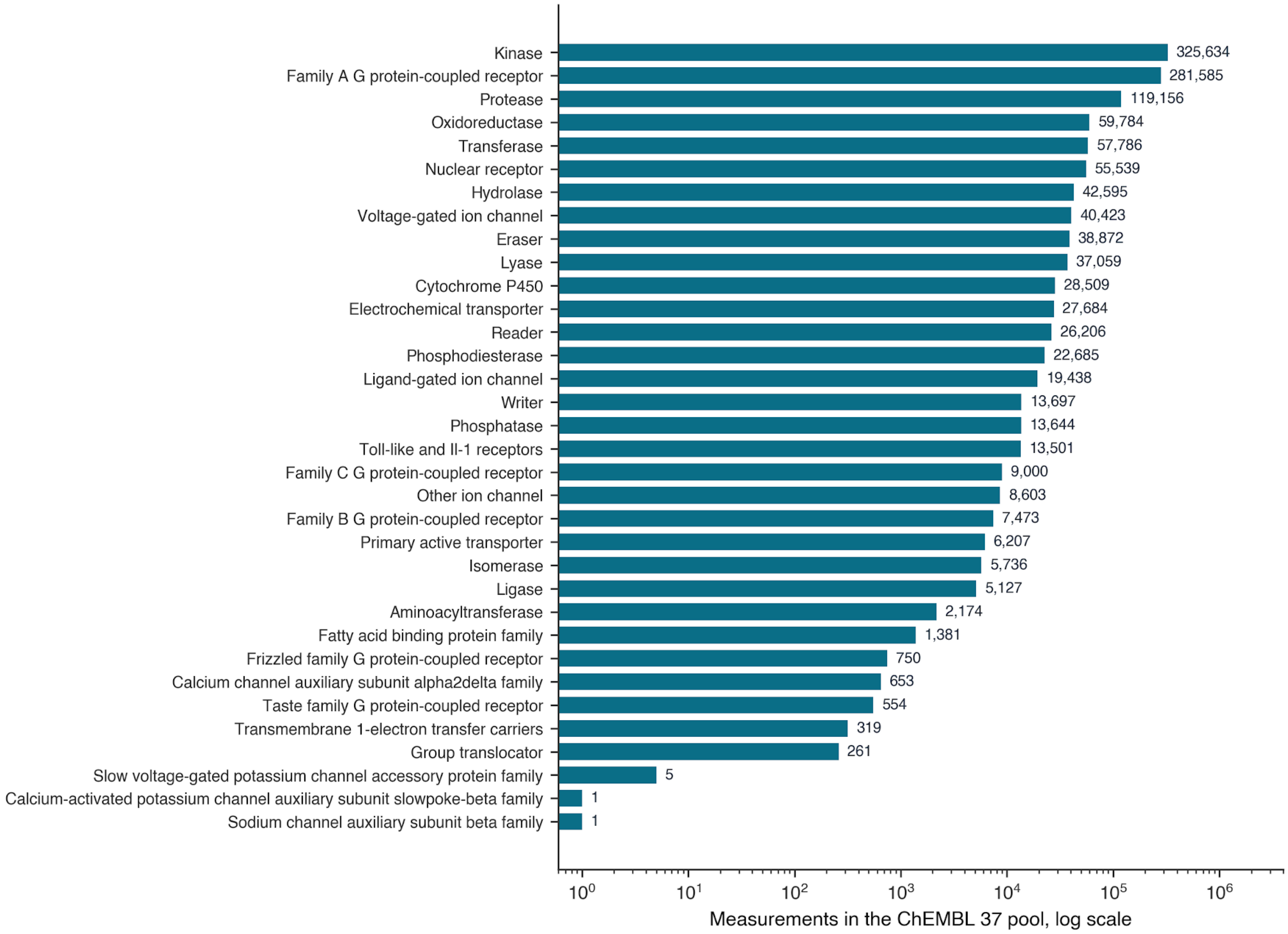


Supplementary Figure S1. The training pool by protein family. The measurement column of Supplementary Table S1 drawn on a logarithmic axis, families ordered by count. Two families hold nearly half the pool between them, and the tail runs down to single readings, which is why the pairings in Supplementary Table S2 are restricted to those carrying 30 or more held-out comparisons.

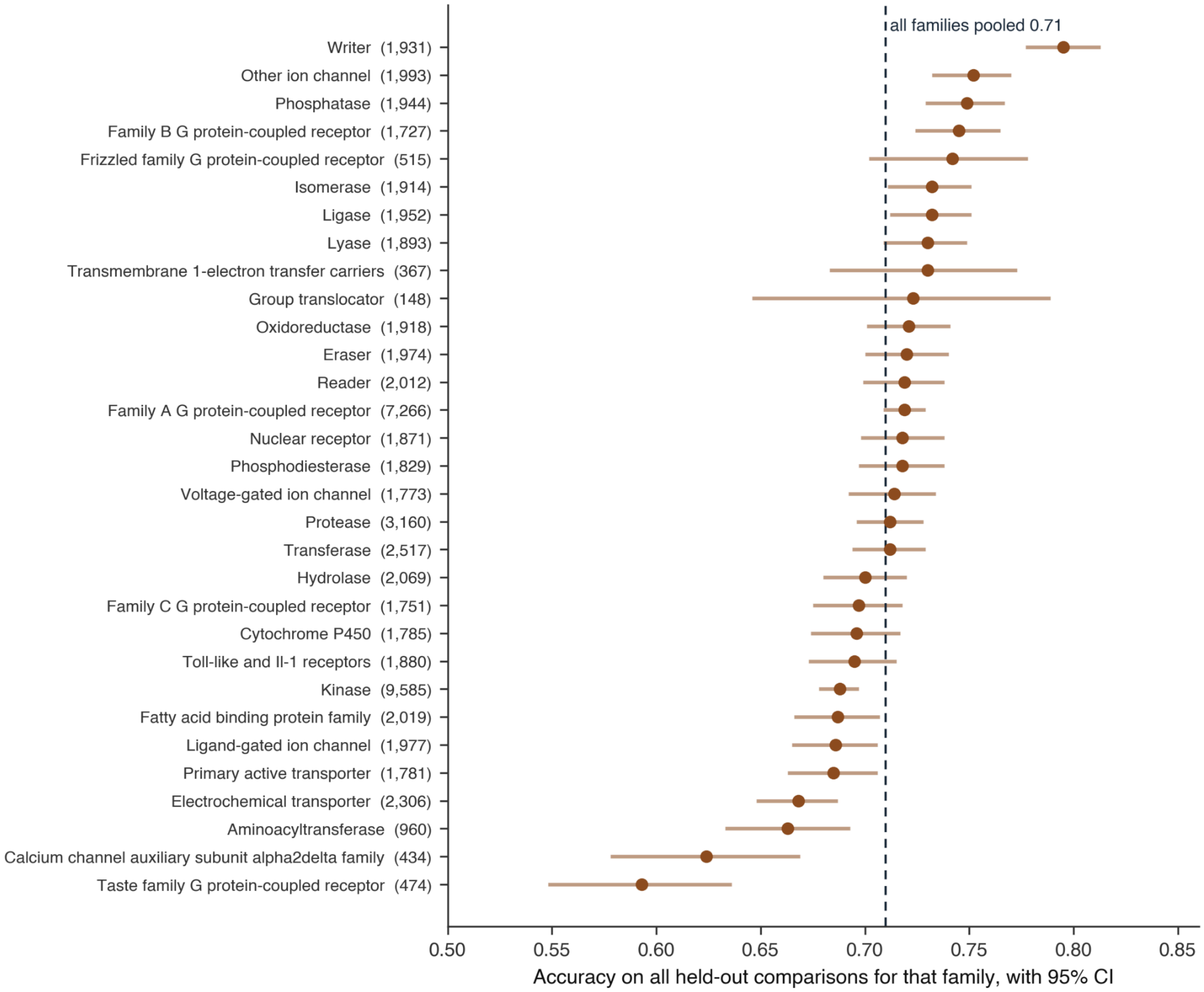


Supplementary Figure S2. Compound-comparison accuracy by family, with 95% confidence intervals. One row per family, ordered by accuracy, over the 31 families carrying held-out compound comparisons, with each family's held-out count in the label. Accuracy is at the ungated operating point, so every held-out comparison for that family is included and nothing is declined. The dashed line is the pooled figure of 0.71 over all 65,725 comparisons. Intervals are wide where a family holds few comparisons, which is the point of showing them; 27 of the 31 families hold out 500 or more. The holdout fraction is set per family precisely so that these rows are comparable in absolute terms, so the ordering here is not a ranking of the families themselves.

Supplementary Table S3. Compound-comparison accuracy by family, at the ungated and 0.80 operating points. The 31 of the compound comparator's 32 roster families that carry held-out comparisons, with 95% confidence intervals, the count behind each figure, and a dash where a family has no comparison reaching strength 0.80. This is the one table in the paper carrying three decimal places, because an interval quoted to two cannot show its own width. The families are ordered by held-out count, which follows from the per-family holdout fraction described in 2.4, independently of pool size.

| Family | Held out | Accuracy, 95% CI | n at 0.80 | Accuracy at 0.80, 95% CI |
|---|---|---|---|---|
| Kinase | 9,585 | 0.688 [0.678, 0.697] | 497 | 0.958 [0.936, 0.972] |
| Family A G protein-coupled receptor | 7,266 | 0.719 [0.709, 0.729] | 485 | 0.930 [0.904, 0.949] |
| Protease | 3,160 | 0.712 [0.696, 0.728] | 276 | 0.953 [0.921, 0.972] |
| Transferase | 2,517 | 0.712 [0.694, 0.729] | 187 | 0.952 [0.911, 0.975] |
| Electrochemical transporter | 2,306 | 0.668 [0.648, 0.687] | 105 | 0.924 [0.857, 0.961] |
| Hydrolase | 2,069 | 0.700 [0.680, 0.720] | 116 | 0.948 [0.892, 0.976] |
| Fatty acid binding protein family | 2,019 | 0.687 [0.666, 0.707] | - | - |
| Reader | 2,012 | 0.719 [0.699, 0.738] | 190 | 0.958 [0.919, 0.979] |
| Other ion channel | 1,993 | 0.752 [0.732, 0.770] | 243 | 0.979 [0.953, 0.991] |
| Ligand-gated ion channel | 1,977 | 0.686 [0.665, 0.706] | 77 | 0.974 [0.910, 0.993] |
| Eraser | 1,974 | 0.720 [0.700, 0.740] | 88 | 0.966 [0.904, 0.988] |
| Ligase | 1,952 | 0.732 [0.712, 0.751] | 95 | 0.968 [0.911, 0.989] |
| Phosphatase | 1,944 | 0.749 [0.729, 0.767] | 118 | 1.000 [0.969, 1.000] |
| Writer | 1,931 | 0.795 [0.777, 0.813] | 174 | 0.989 [0.959, 0.997] |
| Oxidoreductase | 1,918 | 0.721 [0.701, 0.741] | 157 | 0.955 [0.911, 0.978] |
| Isomerase | 1,914 | 0.732 [0.711, 0.751] | 217 | 0.977 [0.947, 0.990] |
| Lyase | 1,893 | 0.730 [0.709, 0.749] | 110 | 0.991 [0.950, 0.998] |
| Toll-like and Il-1 receptors | 1,880 | 0.695 [0.673, 0.715] | 30 | 1.000 [0.886, 1.000] |
| Nuclear receptor | 1,871 | 0.718 [0.698, 0.738] | 89 | 0.989 [0.939, 0.998] |
| Phosphodiesterase | 1,829 | 0.718 [0.697, 0.738] | 76 | 0.987 [0.929, 0.998] |
| Cytochrome P450 | 1,785 | 0.696 [0.674, 0.717] | - | - |
| Primary active transporter | 1,781 | 0.685 [0.663, 0.706] | 75 | 1.000 [0.951, 1.000] |
| Voltage-gated ion channel | 1,773 | 0.714 [0.692, 0.734] | 110 | 0.973 [0.923, 0.991] |
| Family C G protein-coupled receptor | 1,751 | 0.697 [0.675, 0.718] | 83 | 0.988 [0.935, 0.998] |
| Family B G protein-coupled receptor | 1,727 | 0.745 [0.724, 0.765] | 316 | 0.937 [0.904, 0.959] |
| Aminoacyltransferase | 960 | 0.663 [0.633, 0.693] | - | - |
| Frizzled family G protein-coupled receptor | 515 | 0.742 [0.702, 0.778] | - | - |
| Taste family G protein-coupled receptor | 474 | 0.593 [0.548, 0.636] | - | - |
| Calcium channel auxiliary subunit alpha2delta family | 434 | 0.624 [0.578, 0.669] | - | - |
| Transmembrane 1-electron transfer carriers | 367 | 0.730 [0.683, 0.773] | - | - |
| Group translocator | 148 | 0.723 [0.646, 0.789] | - | - |

Supplementary Table S4. Both comparators by endpoint. Accuracy per endpoint on each comparator's own holdout, with the count behind each figure. The asymmetry in the EC50 and Kb rows is the structural difference between the two constructions: an across-family target comparison needs the same endpoint measured at two targets in different families, and agonist data are rarely collected that way. The Kb and EC50 cells for the across-family target comparator are too thin to carry weight and are reported for completeness. The within-family arm carries the agonist endpoints the across-family arm cannot: 2,815 EC50 and 73 Kb comparisons, since two targets of one family are far more often run on the same functional assay than two targets of different ones.

| Endpoint | Target, across families | n | Target, within one family | n | Compound comparator | n |
|---|---|---|---|---|---|---|
| IC50 | 0.78 | 6,577 | 0.80 | 18,533 | 0.71 | 34,481 |
| Ki | 0.70 | 1,440 | 0.78 | 8,825 | 0.72 | 12,900 |
| Kd | 0.57 | 558 | 0.67 | 2,492 | 0.69 | 8,431 |
| EC50 | 0.66 | 114 | 0.81 | 2,815 | 0.71 | 9,375 |
| Kb | - | 0 | 0.78 | 73 | 0.73 | 538 |

Supplementary Table S5. The sibling self-check, and why the three figures are not a ranking. The kinase family's compound-comparison accuracy should land near the Kinase Foundation Model's own within-family result, and Family A G protein-coupled receptor near the GPCR Foundation Model's, which is the strongest single check that the multi-family model is not riding one family. The three splits are not comparable and must not be read as a like-for-like ranking. The Kinase Foundation Model is fitted on the Eidogen-Sertanty Kinase Knowledgebase and tested on ChEMBL, which is held back from its fitting entirely, so its figure is a transfer across corpora rather than a split within one. A compound measured in both records can appear on both sides, which makes its chemistry familiar where ours is withheld, while the corpus it must transfer from is one this model never sees. Neither arrangement is the other's control. The split used here is the strictest of the three, with both compounds unseen and straddling comparisons discarded. The same check on the target axis reads the same way and carries the same warning against ranking the three. Inside Family A G protein-coupled receptor this model reads 0.76 against the GPCR Foundation Model's 0.80, and inside the kinase family 0.73 against the Kinase Foundation Model's 0.75. The kinase figure is the one to read carefully, because that model is fitted on a corpus outside ChEMBL and holds ChEMBL back to test, so its 0.75 measures what a private corpus transfers to the public record while the 0.73 beside it measures what the public record supports on compounds withheld from it. A multi-family model landing within a few points of a single-family model on that family's own question is the result to take from this table, and the three arrangements are why it is not a ranking.

| Model | Headline | n | Split | |
|---|---|---|---|---|
| Kinase Foundation Model, within-family compound comparator | 0.69 | 1,836,100 | not compound-disjoint | |
| GPCR Foundation Model, within-family compound comparator | 0.77 | 762,493 | compound-disjoint, 10%, per target | |
| This work, kinase family | 0.69 | 9,585 | compound-disjoint on both compounds | |
| This work, Family A G protein-coupled receptor | 0.72 | 7,266 | compound-disjoint on both compounds | |
| **Target comparison, within one family** | | | | |
| **Model** | **Accuracy** | **Comparisons** | **Fitted on** | **Tested on** |
| Kinase Foundation Model, within-family target comparator | 0.75 | 3,137,588 | Kinase Knowledgebase, Q2-2026 | ChEMBL, held back entirely |
| GPCR Foundation Model, within-family target comparator | 0.80 | 24,741 | ChEMBL plus Eidogen-curated GPCR data | its own compounds, 10% compound-disjoint holdout stratified across targets |
| This work, within the kinase family | 0.73 | 4,979 | ChEMBL 37 | ChEMBL, compound-disjoint |
| This work, within Family A G protein-coupled receptor | 0.76 | 5,412 | ChEMBL 37 | ChEMBL, compound-disjoint |

Supplementary Table S6. Every classified human target, and the rule that kept or removed it. The 3,213 human single proteins carrying a level-2 ChEMBL class, resolved against the scope rules of 2.1 in the order they are applied. The counts reconcile exactly, 3,213 less 508 leaving the 2,705 of Table 1, with no residual. The four in the last row survived the activity filters and failed elsewhere: three histone methyltransferases whose only exact readings carry the unit string 10^2 uM, which is not on the accepted unit list and whose nanomolar readings are all censored, and one phosphoglycolate phosphatase whose two exact readings belong to compounds with no structure record and therefore no fingerprint. Percentage readings are the largest single cause of loss and they come from the primary literature; high-throughput screening is excluded separately.

| Step | Targets | Rule |
|---|---:|---|
| Carrying a level-2 protein class | 3,213 | |
| No activity row of any kind | 16 | nothing to load |
| Rows exist, none on an accepted endpoint | 362 | endpoint scope, Ki, IC50, Kd, EC50, Kb |
| Endpoint rows exist, all censored or unvalued | 126 | exact readings only |
| Survived the activity filters, still absent | 4 | unit list, or no compound structure |
| **In the pool** | **2,705** | |

Supplementary Table S7. Two families that keep fewer than half their targets. Families losing more than half their human single proteins to the scope rules, counting only families with six or more members so the fraction carries meaning. Both losses are to censored and percentage-only literature, and neither family lost its result with them; the rows therefore remain outside the Discussion.

| Family | Targets kept | Comparisons formed | Accuracy on its own held-out set |
|---|---:|---:|---:|
| Family B G protein-coupled receptor | 15 of 31 | 22,779 | 0.75 |
| Frizzled family G protein-coupled receptor | 4 of 9 | 2,095 | see Supplementary Table S3 |

Supplementary Table S8. Within-family accuracy by family. The 17 of the 30 families carrying within-family comparisons that hold 200 or more of them in the holdout, measured on the 32,738-comparison within-family holdout, with the number of distinct proteins appearing on either side. Two columns govern how a row reads. The proteins column is the first: lyase reaches 0.77 over 15 proteins and phosphodiesterase 0.80 over 22, so those figures describe a handful of well-measured proteins rather than a family. The held-out column is the second: the 13 remaining families hold between 1 and 170 comparisons each and are not scored here. Kinase is the family with the most data and the lowest accuracy of the well-populated rows, at 0.73 over 360 proteins, which is the cost of comparing proteins that resemble one another.

| Family | Held out | Accuracy | Proteins |
|---|---|---|---|
| Family A G protein-coupled receptor | 5,412 | 0.76 | 160 |
| Kinase | 4,979 | 0.73 | 360 |
| Protease | 3,889 | 0.81 | 134 |
| Lyase | 2,792 | 0.77 | 15 |
| Eraser | 2,315 | 0.79 | 36 |
| Transferase | 1,993 | 0.85 | 90 |
| Cytochrome P450 | 1,755 | 0.76 | 28 |
| Nuclear receptor | 1,601 | 0.81 | 33 |
| Electrochemical transporter | 1,207 | 0.79 | 45 |
| Oxidoreductase | 1,137 | 0.79 | 86 |
| Phosphodiesterase | 938 | 0.80 | 22 |
| Reader | 924 | 0.83 | 53 |
| Toll-like and Il-1 receptors | 922 | 0.94 | 6 |
| Voltage-gated ion channel | 811 | 0.81 | 48 |
| Hydrolase | 600 | 0.77 | 70 |
| Phosphatase | 575 | 0.74 | 40 |
| Ligand-gated ion channel | 250 | 0.69 | 31 |

Supplementary Table S9. The comparisons the record allows. Available counts every endpoint-matched pair of readings on one compound that is not a tie, before the caps of 2.2 are applied; formed is what those caps keep, 50 per compound in each arm and 20,000 per family pairing and endpoint. Kinase against kinase is 1,127,614 of the available within-family total, 69% of it, which is why a within-family result is reported by family rather than pooled. The caps, not the record, set the size of the fitted sets here, and the across-family arm is the one the record itself bounds (Figure 3).

| Comparison | Available | Formed after caps | Held out |
|---|---|---|---|
| Target, across families | 518,194 | 89,888 | 8,689 |
| Target, within one family | 1,643,762 | 333,893 | 32,738 |
| Compound preference | not capped this way | 2,519,433 | 65,725 |
